# The non-linear development of the right hemispheric specialization for human face perception

**DOI:** 10.1101/122002

**Authors:** Aliette Lochy, Adelaïde de Heering, Bruno Rossion

## Abstract

The developmental origins of human adults’ right hemispheric specialization for face perception remain unclear. On the one hand, infant studies have generally shown a right hemispheric advantage for face perception. On the other hand, the adult right hemispheric lateralization for face perception is thought to slowly emerge during childhood, due to reading acquisition, which increases left lateralized posterior responses to competing written material (i.e., visual letters and words). Since methodological approaches used in infant and children usually differ, resolving this issue has been difficult. Here we tested 5-year-old preschoolers varying in their level of visual letter knowledge with the same fast periodic visual stimulation (FPVS) paradigm leading to strongly right lateralized electrophysiological occipito-temporal face-selective responses in 4- to 6-month-old infants (de Heering & Rossion, 2015). Children’s face-selective response was much larger and more complex than in infants, but did not differ across hemispheres. However, there was a small positive correlation between preschoolers’ letter knowledge and their right hemispheric specialization for faces. These observations suggest that several factors contribute to the adult right hemispheric lateralization for faces, and point to the value of FPVS coupled with electroencephalography to assess specialized face perception processes throughout development with the same methodology.

## Introduction

Hemispheric lateralization of brain function is well established in humans as well as in other animal species. Yet the reasons for this lateralization are still largely unknown and debated (Corballis, 2009; Davidson & Hugdahl, 1995; Güntürkün et al., 2000). In humans, the right hemisphere (RH) is dominant in the perception of faces of conspecifics, a critical brain function for social interactions. This RH dominance for face perception has been initially supported by lesion studies, showing that a right ventral occipito-temporal lesion, typically associated with left upper visual field defects, is both necessary and sufficient to cause prosopagnosia, i.e. a severe and sometimes specific impairment at individual face recognition (Hecaen & Angelergues, 1962; Meadows, 1974; Sergent & Signoret, 1992; Davies-Thompson, Pancaroglu, & Barton, 2014; Rossion, 2014 for reviews) Divided visual field studies and chimeric face effects have also pointed to a right hemisphere advantage in face perception (Gilbert & Bakan, 1973; Hillger & Koenig, 1991; Kolb, Milner, & Taylor, 1983; Levy, Trevarthen, & Sperry, 1972; Rizzolatti, Umiltà, & Berlucchi, 1971), a conclusion largely corroborated over the past two decades by numerous neuroimaging studies (Frässle, Paulus, Krach, & Jansen, 2016; Kanwisher, McDermott, & Chun, 1997; Rossion, Hanseeuw, & Dricot, 2012; Sergent, Ohta, & MacDonald, 1992) and high-density electroencephalographic (EEG) recordings on the human scalp (the right lateralized N170 potential evoked by faces; e.g., (Bentin, Allison, Puce, Perez, & McCarthy, 1996; Rossion & Jacques, 2011) for review). More recently, a strong right hemispheric dominance for face-selective responses in the human ventral occipito-temporal cortex (VOTC) has also been reported with intracerebral electrophysiological recordings (Jonas et al., 2016).

Since the critical brain regions involved in face perception are right lateralized in human adults, understanding *when* this right hemispheric lateralization emerges during human development and *which factors* drive this specialization is important to deepen our understanding of human face perception.

de Schonen and Mathivet (1989) initially proposed that the right hemispheric specialization for face perception emerges relatively *early* during development, i.e. already at a few months of age. Their proposal was based on the observation that 4- to 9-month-old infants saccade faster towards the picture of their mother’s face than the matched picture of a stranger’s face when these pictures are presented in the left visual field (LVF) but not in the right visual field (RVF; de Schonen & Mathivet, 1990; de Schonen, Gil de Diaz, & Mathivet, 1986). Along the same line, right hemisphere but not left hemisphere early deprivation of visual input for several months (between 6 weeks and 3 years) impairs the development of the adult expert (i.e., holistic/configural) face processing system (Le Grand, Mondloch, Maurer, & Brent, 2003). At the neural level, studies using functional near-infrared spectroscopy (fNIRS) have also generally shown a significant RH advantage for faces over control visual stimuli in 5- to 8-month-old infants (e.g., Otsuka et al., 2007; see Otsuka, 2014 for review). Most recently, de Heering and Rossion (2015) exposed 4- to 6-month-old infants to numerous and highly variable natural images of faces inserted periodically (1 out of 5) in a fast (6 Hz) stream of non-face object images while recording their electroencephalogram (EEG) (Figure 1A). Strikingly, the face-specific response at the frequency at which faces were presented in the sequence (1.2 Hz = 6Hz/5) was localized almost exclusively over the infants’s right occipito-temporal cortex (Figure 1B). Importantly, the specific response to faces recorded over the right occipito-temporal cortex was not found for phase-scrambled images, ruling out potential low-level visual accounts of the effect (de Heering & Rossion, 2015).

Altogether, these observations support de Schonen & Mathivet, 1989’s hypothesis that the right hemisphere takes precedence over the left hemisphere, which might be the result of its faster maturation rate at the time at which the infants’ visual system mainly extract low spatial frequencies, and therefore global information, from facial inputs (Sergent, 1982)

However, according to a recent hypothesis, the right hemispheric specialization for face perception would rather emerge relatively *late* during development, i.e. when children learn to read, and would gradually increase through adolescence (Behrmann & Plaut, 2015; Dehaene, Cohen, Morais, & Kolinsky, 2015; Dundas, Plaut, & Behrmann, 2012). This hypothesis generally rests on the general view that right hemispheric specialization for faces – and of other functions such as spatial perception – follows the left lateralization for language functions (Corballis, 1991; Lhermitte, Chain, Escourolle, Ducarne, & Pillon, 1972). More specifically, it states that the right hemisphere becomes dominant for face perception due to the gradual specialization of the left VOTC after children’s exposure to visual words during reading acquisition. This specialization would then compete with the representation of faces in the left hemisphere, resulting in face representations mainly located in the RH (Behrmann & Plaut, 2015). This view is supported by several findings. First, the behavioral left visual field advantage caused by the RH superiority for face processing correlates positively with reading competence in school children and young adolescent (Dundas et al., 2012). Moreover and contrary to adults, children up to 9-12 years of age do not systematically show a larger face-evoked N170 for faces in the RH than in the LH (Dundas, Plaut, & Behrmann, 2014; Itier & Taylor, 2004b; Kuefner, de Heering, Jacques, Palmero-Soler, & Rossion, 2010; Taylor, McCarthy, Saliba, & Degiovanni, 1999). Third, literacy changes the hemispheric balance of neural response to faces in both and adults and 10-year-old children, with a slight decrease of neural activity in the left fusiform gyrus and a clearer increase in the homologous area of the right fusiform gyrus (Dehaene et al., 2010; Monzalvo, Fluss, Billard, Dehaene, & Dehaene-Lambertz, 2012; respectively). Finally, left-handed individuals, who as a group show greater variability with respect to hemispheric language dominance than right-handed individuals, also show greater variability in their degree of RH lateralization of faces as evidenced from both behavioral (Dundas, Plaut, & Behrmann, 2015) and neural measurements (fMRI: Bukowski, Dricot, Hanseeuw, & Rossion, 2013; Frässle et al., 2016; EEG: Dundas et al., 2015).

In sum, there is a striking contrast between evidence collected in young infants, supporting the early emergence of a RH lateralization for face perception independent of reading acquisition, and evidence gathered from children, adolescents and adults, rather favoring a late and gradual RH lateralization of face perception emerging as a consequence of reading acquisition and reading skills.

So far, these views, based on different sets of evidence, have been hard to reconcile. The major reason for this difficulty is that studies in infants and children have been conducted with different techniques and paradigms. For instance, while fNIRS, visual field dominance paradigms and visual preference/adaptation paradigms have been generally used to test infants, fMRI and explicit behavioral tasks in divided visual field presentations have rather been used with children. Second, even with a technique readily applicable in both infants and children/adults such as the recording of event-related potentials (ERPs) in EEG, responses remain difficult to compare across populations. For instance, the occipitotemporal N170 found in adults and also children from 5 years of age (Itier & Taylor, 2004a; Kuefner et al., 2010), is not observed in young infants, in which ERP components to faces peak later and over medial occipital sites, without lateralization (i.e., the N290, de Haan & Nelson, 1999; de Haan, Pascalis, & Johnson, 2002; Gliga & Dehaene-Lambertz, 2007; Hoehl & Peykarjou, 2012).

The primary goal of the present study was to start filling this gap in our knowledge by applying the exact same paradigm of de Heering and Rossion (2015) showing right lateralized face-selective responses in 4- to 6-month-old infants to a population of preschool children varying in their knowledge of letters and already showing a left lateralized specialization for this kind of material.

At many levels, this Fast Periodic Visual Stimulation (FPVS; Rossion, 2014b) approach combined with EEG appears ideally positioned to shed light on the origin and the developmental course of face perception and of its RH lateralization. First, FPVS leads to the recording of high signal-to-noise ratio brain responses usually termed “Steady-State Visual Evoked Potentials” (SSVEPs, after (Regan, 1989; Regan, 1966); see Norcia, Appelbaum, Ales, Cottereau, & Rossion, 2015 for review), which can be identified objectively in the frequency domain after Fourier Transform. Hence, it has been used in classical infant studies to inform about early development of primary visual functions (e.g., Braddick, Wattam-Bell, & Atkinson, 1986; Norcia, Tyler, & Hamer, 1988). Second, it does not require any explicit task that might be contaminated by motivational or decisional factors that are particularly hard to control in young individuals, and differ greatly across development. Third, it leads to electrophysiological responses that can be directly quantified at the individual level, allowing to test relationship between these responses and behavioral measures in children for instance (e.g., Lochy, Van Reybroeck, & Rossion, 2016). Finally, and more specifically, this approach has been successfully extended to capture high level generic face categorization responses in adults (e.g., Liu-Shuang, Norcia, & Rossion, 2014; Rossion et al., 2015).

With this approach in hand, the first goal of the present study was to test whether the right hemispheric lateralization of face-selective responses observed in 4-6 months old infants is also observed in preschool children of 5-6 years of age. We targeted specifically this children population because they have not been exposed to formal reading acquisition, yet they show variable knowledge of visual letters, this knowledge driving their early left hemispheric lateralization for this material in FPVS-EEG (Lochy et al., 2016). Since the current group of preschoolers has also been tested in a previous study with orthographic material (Lochy et al., 2016), the second goal was to examine the potential relationship between letter knowledge, left lateralization of the response to letters, and lateralization of the response to faces.

## Material and Methods

### Participants

The data of thirty-four children (20 males, mean age = 5.51 years; range = 5.01-5.94 years), with normal/corrected-to-normal vision, was collected after the parents gave informed consent for a study approved by the Biomedical Ethical Committee of the University of Louvain (Belgium). Children were unaware of the goal of the experiment and that a change of stimulus type occurred at a periodic rate during stimulation. These children were part of a larger sample tested in a visual letter discrimination experiment that revealed a specific response for letters (words or pseudo-words) among pseudo-letters in the left hemisphere (O1) that also correlated with their letter knowledge (Lochy et al., 2016). They underwent a screening battery with sub-tests of the WISC-R, visuo-attentional capacities, vocabulary, and reading competencies.

### Stimuli

Two-hundred and fifty images of various objects (animals, plants, man-made objects) and 50 images of faces, collected from the internet, were used, as in previous studies (de Heering & Rossion, 2015; Rossion et al., 2015). They differed in terms of viewpoint, lighting conditions and background (Fig.1A). They were all resized to 200 × 200 pixels, equalized in terms of luminance and contrast in Matlab (Mathworks), and shown in the center of the screen at a 800 × 600 pixel resolution. At a testing distance of 40 cm, they subtended approximately 13 by 13 degrees of visual angle.

**Figure 1.**
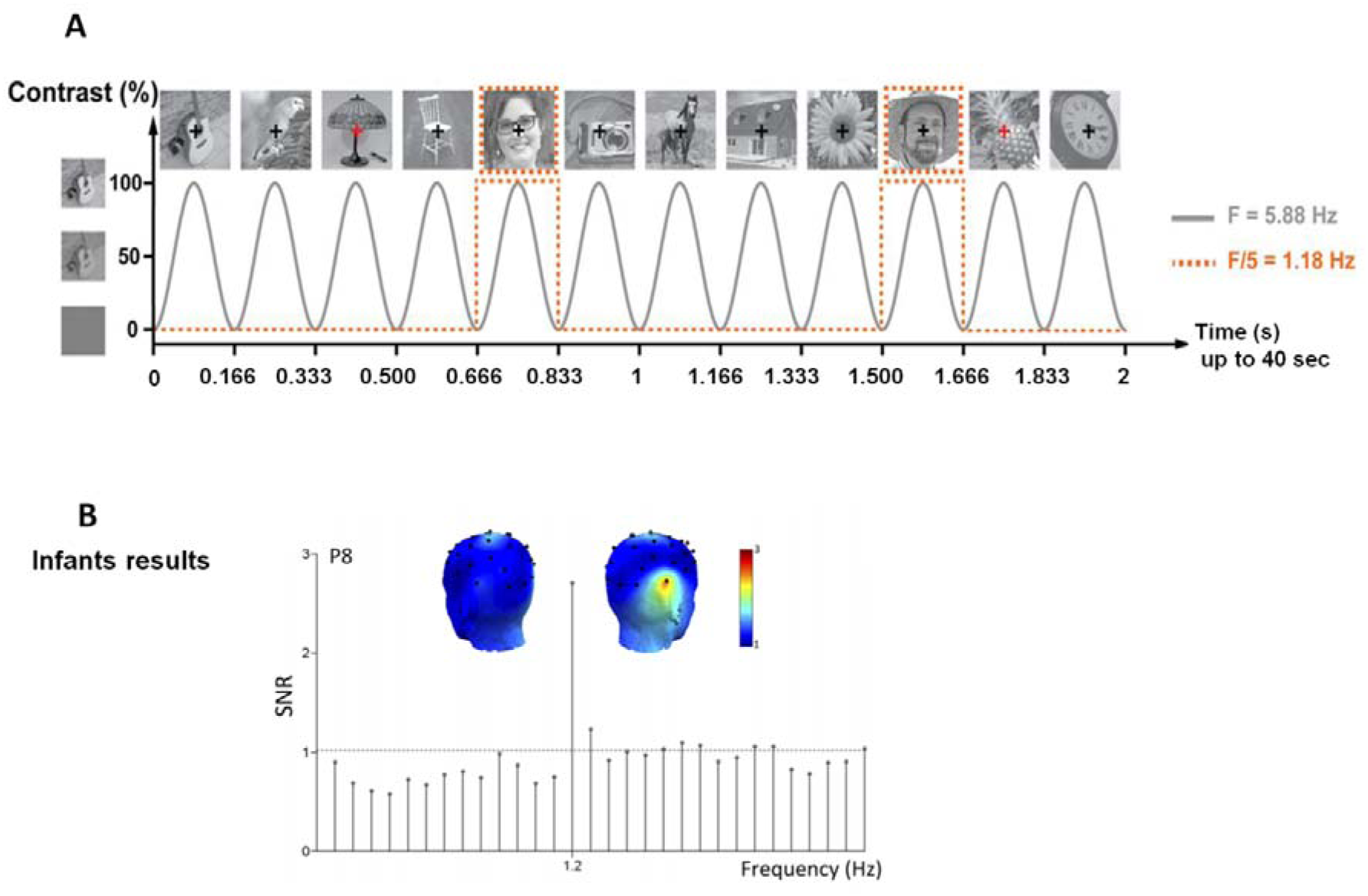
***A.*** A visual stimulation sequence where base stimuli are constituted of various non-face objects, in which highly variable images of faces (various identities, viewpoints, …) are inserted periodically (i.e., every 5 stimuli) (from de Heering & Rossion, 2015; Rossion et al., 2015). Stimulation mode: 6 stimuli were presented per second with a sinusoidal contrast modulation, and every fifth item was a face stimulus. Each child viewed 2 40-second stimulation sequences (i.e., 47 faces inserted in non-face objects). ***B.*** Results (Grand Averaged frequency spectrum in signal-to-noise ratio (SNR) obtained in 4- to 6-month-old infants with the same paradigm by de Heering and Rossion (2015), showing a clear peak of activation at 1.2Hz located on the right occipito-temporal lobe, reflecting generic face categorization.

### Procedure

Stimulation was virtually identical to the study of de Heering and Rossion (2015) in infants, except for the duration of stimulation sequences (20 seconds in infants) and their number (variable in individual infants). Each stimulation sequence started with a fixation cross displayed for 2 – 5 seconds, 2 seconds of gradual stimulation fade-in, 40 seconds of stimulation sequence, and 2 seconds of gradual fade-out. Stimuli were presented through sinusoidal contrast modulation at a base frequency rate of 6Hz (i.e., one item every 166.66 ms, hence each item reached full contrast after 83 ms) (Fig. 1A). Given that the stimulus can be recognized at very low contrast (i.e., 20% or less), the actual duration of stimulus visibility approximates 140 ms.

Every sequence followed the same structure: base stimuli (non-face objects) (B) were presented at 6Hz, and every fifth item was a deviant stimulus of the category of interest, i.e., faces (1.2Hz, thus every 833 ms) such as: OOOOFOOOOFOOOOF…. (Fig. 1A). MATLAB 7.8 (The Mathworks) with PsychToolbox ((Brainard, 1997) see http://psychtoolbox.org/) was used for stimulus display. Since the approach provides a very high signal-to-noise ratio (SNR), even in young infants (de Heering & Rossion, 2015), only two stimulation sequences of 40 seconds were used, and were initiated manually to ensure low-artifact EEG signals. During stimulation, children fixated a central cross and were instructed to press the space bar for any brief (200 ms) color change of the fixation cross (blue to red; 6 changes randomly timed per sequence). The goal of this orthogonal task was to maintain their level of attention as constant as possible throughout the experiment. Children performed this task almost at ceiling (mean: 90%; SE: 0.02), showing high attention to the stimuli presented on the screen, with average response times of 733ms (SE: 17.08).

### EEG acquisition and preprocessing

Children were seated comfortably at 1 m from the computer screen in a quiet room of the school. EEG was acquired at 1024Hz using a 32-channel Biosemi Active II system (Biosemi, Amsterdam, Netherlands), with electrodes including standard 10-20 system locations (http://www.biosemi.com). The magnitude of the offset of all electrodes, referenced to the common mode sense (CMS), was held below 50 mV. All EEG analyses were carried out using *Letswave* 5 (http://nocions.webnode.com/letswave), and *Matlab 2012* (The Mathworks) and were virtually identical to analyses performed on the same type of data collected in adults (Rossion et al., 2015) and infants (de Heering & Rossion, 2015). After FFT band-pass filtering around 0.1 and 100Hz, EEG data were segmented to include 2 seconds before and after each sequence, resulting in 44-second segments (−2 – 42 s). Data files were then resampled to 250Hz to reduce file size and data processing time. Artifact-ridden or noisy channels were replaced using linear interpolation (no more than two electrodes for each participant). All channels were re-referenced to the common average. EEG recordings were then segmented again from stimulation onset until 39.996 seconds, corresponding exactly to 48 complete 1.2Hz cycles within stimulation. This corresponds to the largest amount of complete cycles of 833 ms at the face stimulation frequency (1.2Hz) within the 40 seconds of stimulation period.

### Frequency domain analysis

The two trials were averaged in the time domain for each individual participant, in order to increase SNR. A Fast Fourier Transform (FFT) was applied to the averaged time-window, and normalized amplitude spectra were extracted for all channels. This yielded EEG spectra with a high frequency resolution (1/39.996 s = 0.025Hz), increasing SNR (Rossion, 2014c) and allowing unambiguous identification of the response at the exact frequencies of interest (i.e., 6Hz for the base stimulation rate and 1.2Hz and its harmonics for the face stimulation). To estimate SNR across the EEG spectrum, amplitude at each frequency bin was divided by the average amplitude of 20 surrounding bins (10 on each side) (Rossion, Prieto, Boremanse, Kuefner, & Van Belle, 2012). To quantify the responses of interest in microvolts, the average voltage amplitude of the 20 surrounding bins (i.e., the noise) was subtracted out (Retter & Rossion, 2016). Based on the grand-averaged amplitude spectrum, Z-scores were computed at every channel in order to assess the significance of the response at the face stimulation frequency and harmonics, and at the base rate and harmonics (Lochy et al., 2015; Liu-Shuang, Norcia, Rossion, 2014). Z-Scores larger than 1.64 (p < 0.05, one-tailed, signal > noise) were considered significant. The face-selective response was significant up to 10.8Hz (9 harmonics). In order to assess the significance of responses in individual participants, Z-scores were computed on the sums of those harmonics significant at the group-level (from 1.2Hz to 10.8Hz excluding the base rate) and considered significant if larger than 1.64. Finally, to quantify the periodic response distributed on several harmonics, the baseline subtracted amplitudes of significant harmonics (excluding the base stimulation frequency) were summed for each participant (Retter & Rossion, 2016).

## Results

Scalp topographies and EEG spectra of grand-averaged data showed a clear (i.e., SNR >2, >100% increase of signal; sum of baseline-corrected amplitudes > 4μV) face-selective response at 1.2Hz as well as at its harmonics (2.4 Hz, etc.) on posterior electrodes (Fig.2). Strikingly, and contrary to infants (de Heering & Rossion, 2015; see Figure 1A) and adults (Rossion et al., 2015), preschoolers did not show any RH lateralized brain response (Fig.2).

The sum of baseline corrected amplitudes was computed on 9 harmonics excluding the base rate (1.2Hz to 10.8Hz) as determined by grand-averaged data (F/5 to 9F/5) (maximal number of consecutive harmonics with a Z-score >1.64). The largest responses were recorded at electrodes PO3-PO4, O1-O2, and P7-P8 (the left electrodes PO3, P7 and O1 and the right electrodes PO4, P8, and O2 are grouped into left and right ROI for purpose of display in Fig. 2). A 2x3 repeated measures ANOVA with *Hemisphere* (left, right) and *Electrode Site* (posterior -O1/O2-, lateral -P7/P8-, and dorsal -PO3/PO4-) as within-subject factors was conducted on participants’ sum of baseline corrected amplitudes. There was a significant effect of *Electrode Site* [F(2,66)=4.938; p<0.01], with responses being stronger at dorsal sites (PO3/PO4, 4.011μV, SE0.308) than at lateral or posterior sites (P7/P8: 2.9μV, SE=0.305; and O1/O2: 3.423μV, SE=0.265). There was no effect of *Hemisphere* [F(1,33)<1] (3.428μV, SE=0.226 and 3.461μV SE =0.291 for LH and RH respectively) and no interaction between the two factors [F(2,66)=2.750; p=0.08]. Crucially we did not find any significant difference between the 2 hemispheres at any of the electrode sites (dorsal, lateral, posterior) considered into the analyses (all p >0.15; O2-O1=0.065μV; P8-P7=0.639μV; PO4-PO3= −0.607μV). Paired comparisons confirmed that brain responses were significantly stronger at dorsal sites (PO3/PO4) than at lateral (P7/P8) (p<0.007) or posterior (O1/O2) (p<0.04) sites. Lateral and posterior sites did not differ (p=0.18).

Importantly, the face-selective response was significant in every individual child tested on at least two electrodes in these ROIs, as revealed by Z-scores (p<0.05). Almost half of them (N=16) had a significant response on all 6 posterior channels. Even with a more stringent statistical criterion (see Fig) of p<0.001 (Z>3.1), more than half of the participants still displayed a significant response on most channels, and 28 out of 35 on PO3.

**Figure 2.**
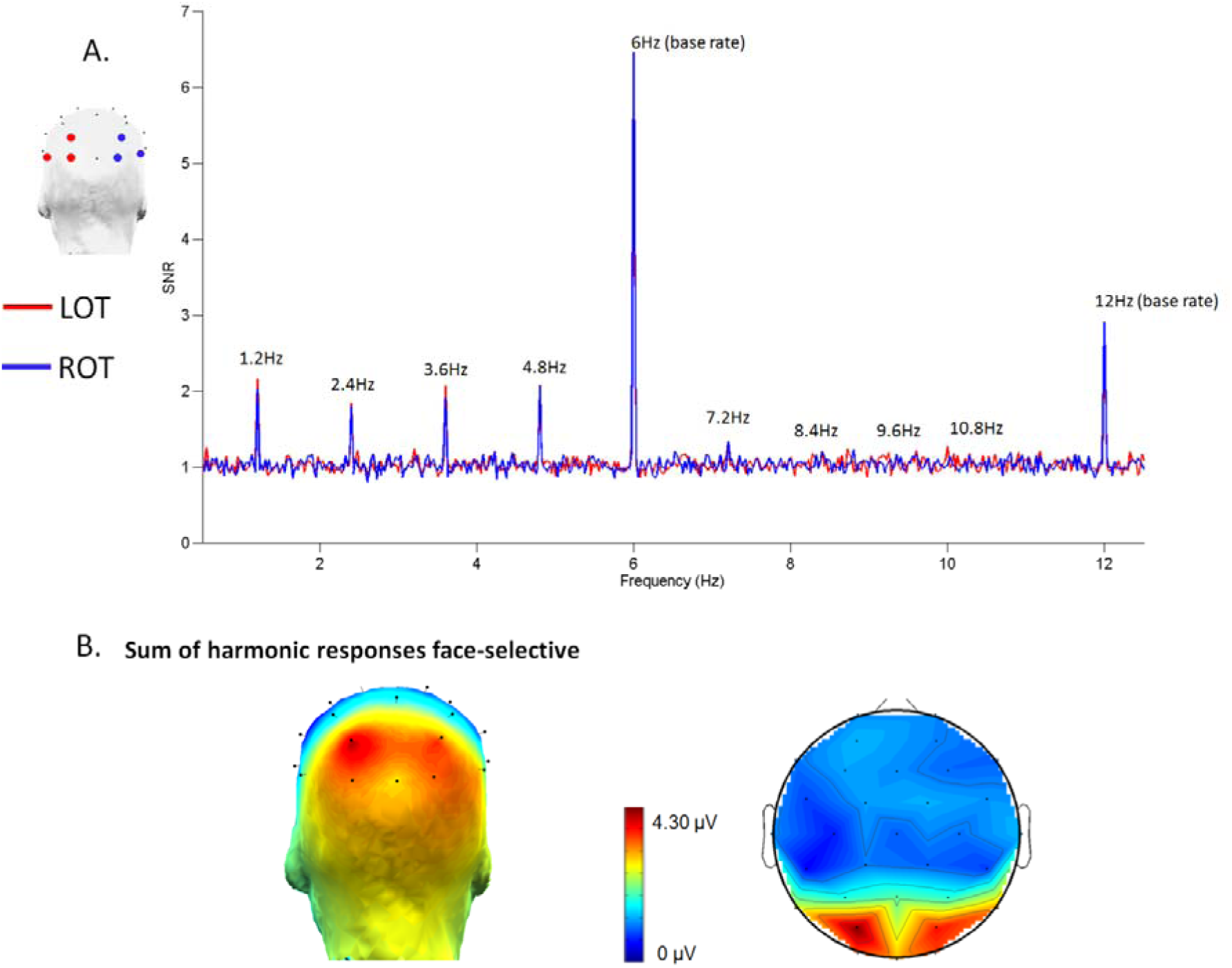
SNR response spectra and scalp topographies (sum of harmonics amplitude in microvolts, see Retter & Rossion, 2016) of the brain responses in the face categorization task. ***A*.** SNR response spectra for left and right ROI, showing significant responses at the F/5 (1.2Hz) frequency and its harmonics, as well as at the base stimulation frequency (6Hz) and harmonics (12 Hz, etc.), common for objects and faces. ***B*.** Average topographical map for the face-selective responses based on the sum of corrected amplitudes at significant harmonics: back view (left) and 2D flat map (right).

We further assessed the potential relationship between lateralization of the face-response with preschoolers’ behavioral measures of reading (production and recognition of grapheme-phoneme correspondences), as well as with the lateralization of their brain responses to letters (extracted from Lochy et al., 2016; where the same paradigm was used with base stimuli at 6Hz stimulation frequency constituted of pseudo-letters and deviant stimuli at 1.2Hz stimulation frequency constituted of letter-strings). To this end, we first computed lateralization scores (LS) by subtracting the amplitude values of the left hemisphere (LH) from values in the right hemisphere (RH) (RH-LH) while focusing on 2 regions-of-interest - the left occipito-temporal cortex (LOT = O1, P7, PO3) and the right occipito-temporal cortex (ROT = O2, P8, PO4) – and then 2 electrodes-of-interest located at the dorsal site: PO3/PO4, associated with the strongest response. Positive and negative values would respectively indicate a right and a left lateralization.

Next we correlated these scores to preschoolers’ behavioral measures (Fig. 3A). Assessing our main hypothesis, we found that children’s letter recognition scores were modestly but significantly correlated with their lateralization score on the dorsal electrode site PO3/PO4 showing the largest response (Spearman Rho=0.296; p=0.045), with a nonsignificant trend on the ROI lateralization score (ROT-LOT) (Spearman Rho=0.256; p=0.072). That is, the right lateralization for faces increased with the number of letters recognized by the children. However, letter production did not correlate with the lateralization score for faces on PO3/PO4 (rho=0.179; p=0.156) or on the ROI (ROT-LOT) (rho=0.175; p=0.162). Note that these two behavioral measures correlated negatively with the lateralization score for letters on O1/O2 (production: rho=−0.335; p=0.026; and recognition, rho=−0.428; p=0.006, Fig. 3B), where the response to letters is maximal on O1 and was used as the letter discrimination response in (Lochy et al., 2016). This replication of a significant correlation with another analysis approach, i.e., when using a lateralization score rather than the raw amplitude in the LH as in (Lochy et al., 2016), confirms that the more letters are known, and the more the LH responds to letter-strings.

Finally we assessed whether the brain responses to letters and to faces were negatively related, that is, whether the more left lateralized the response to letters and the more right lateralized the response to faces. However, there was no significant correlation between lateralization scores in the two types of discriminations (letters vs. faces) whether on ROIs (ROT-LOT) (Spearman Rho=−0.166; p=0.174) or on electrodes with largest responses in each type of discrimination (O2-O1 for letters and PO4-PO3 for faces) (Spearman Rho=−0.122; p=0.245).

**Figure 3.**
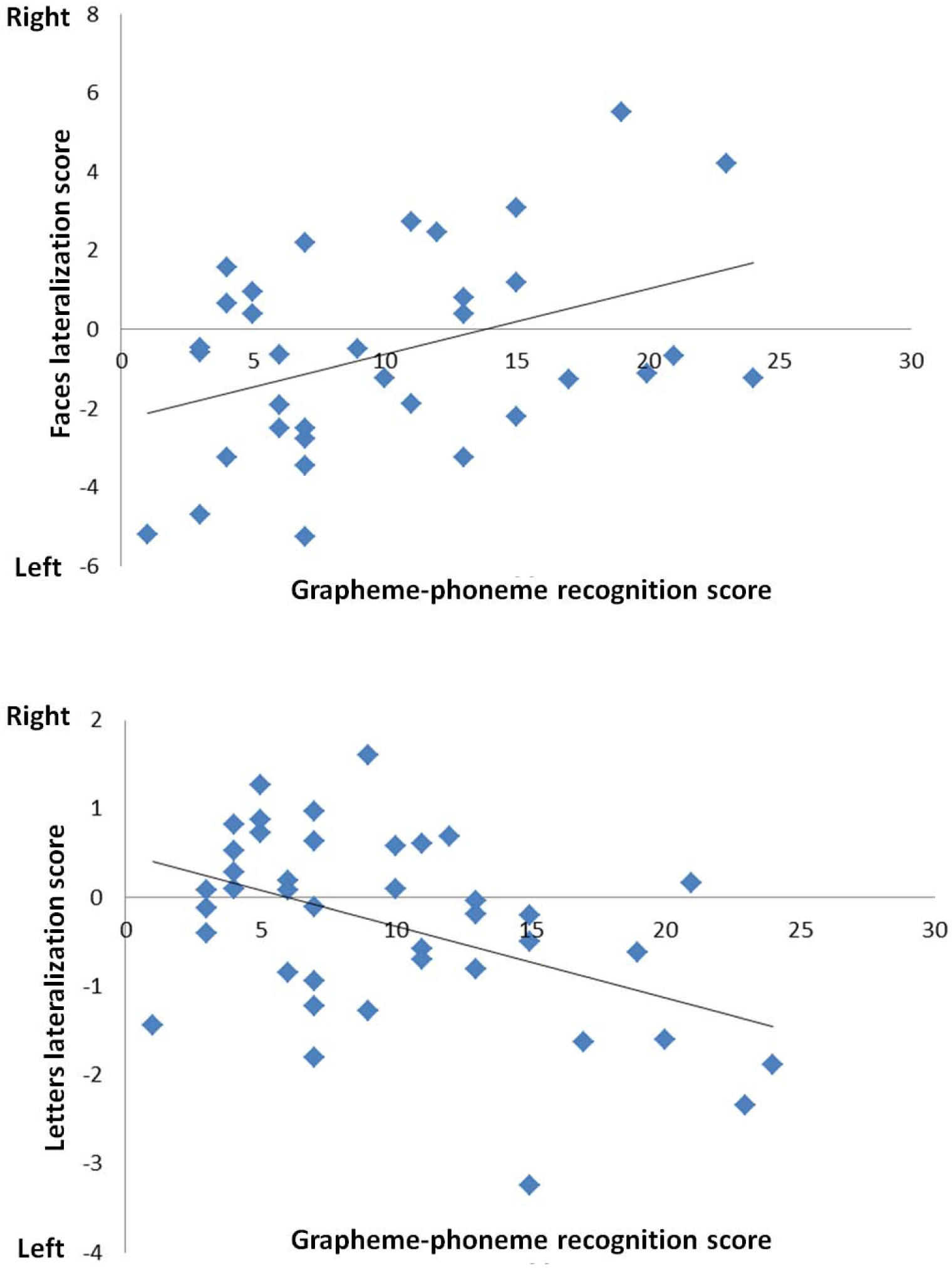
Scatter plots of correlations between a behavioral reading task (grapheme-phoneme recognition, GP) and lateralization scores. ***A*.** Positive relationship (rho=0.29) between GP recognition and RH lateralization score for faces (amplitude at PO4 (RH)-amplitude at PO3 (LH)). ***B*.** Negative relationship (rho= −0.43) between GP recognition and RH lateralization score for letters (amplitude at O2 (RH) – amplitude at O1 (LH)).

## Discussion

Thanks to a FPVS-EEG paradigm used previously in adults (Liu-Shuang, Torfs, & Rossion, 2016; Rossion et al., 2015) and infants (de Heering & Rossion, 2015), we found a robust face-selective electrophysiological response in 5-year-old preschoolers. This face-selective, or face categorization, response was obtained in a few minutes of recording only and was significant in all individual participants on at least two electrodes in the regions of interest. As noted in these previous studies, this response reflects a real high-level perceptual categorization process: while the common response to faces and objects project to a 6 Hz base rate response the 1.2 Hz frequency response and its harmonics appears only if children *discriminate* face images from a wide variety of nonface objects and, in order to maintain periodicity, if they *generalize* this response across the majority of the widely variable face images presented. Moreover, the face categorization response identified in this paradigm cannot be accounted for by low-level visual cues such as differences in amplitude spectrum between faces and objects (see (Rossion et al., 2015).

Five-year-old children’s robust face categorization response may not appear surprising, given that preschool children are already able to perform complex object categorizations in explicit behavioural tasks (e.g., (Bornstein & Arterberry, 2010). However, there is, to our knowledge, very little data on generic face categorization (often referred to as “face detection”) in children. Carbon et al. (Carbon, Grüter, & Grüter, 2013) found that explicit detection of faces in ambiguous (two-tone) Mooney figures was almost adult-like in young children (95% beginning with the age group 6-10; 86% in 2-5 year old), but this is a very different task than what is evaluated in the present study. Here, our results show that five-year-old children are able to categorize widely variable images of faces *automatically*, i.e. without explicit instruction to do so, and extremely *rapidly*, i.e. at a glance: each facial image appears for about 140 ms in this paradigm (Fig 1A), and is forward- and backward-masked by nonface objects.

Nevertheless, since the very same paradigm already gave clear face categorization responses in 4-6 months old infants (de Heering & Rossion, 2015), we fully expected the present observation of a robust FC response in 5-year-old children. The unexpected result, from our point of view at least, is that the face categorization response obtained with this paradigm in 5-year-old children is not at all right lateralized, contrary to the observation made on infants (de Heering & Rossion, 2015)as well as in the majority of typical adults (Liu-Shuang et al., 2016; Retter & Rossion, 2016; Rossion et al., 2015). Besides the bilateral face response at the group level, we found a positive relationship between the number of letters known individually by children, as assessed by a letter recognition task, and their right lateralization for faces as assessed with a lateralization score. There was, however, no relationship between the degree of their left lateralization to words and the right lateralization to faces.

Thus, altogether, these observations suggest that there is a non-linear developmental trajectory in the RH lateralization for generic categorization of faces. The initial right lateralized face-selective response observed in 4-6 months old infants with the exact same paradigm (de Heering & Rossion, 2015) becomes bilateral between a few months and a few years of age, i.e. before formal reading acquisition. Then, between the beginning of school at age 5-6 and adulthood, further changes which may be partly related to visual language acquisition occur, resulting in a right-lateralized face-selective response at adulthood (Rossion et al., 2015). In addition, between infancy and childhood, the face categorization response does not only become much larger and complex as in adults (i.e., distributed over many harmonics while it was restricted to the first harmonic in infants), but it is more widely and dorsally distributed compared to the ventral occipito-temporal response found in infants and adults. These observations are discussed further below.

### Bilateralization of face-selective responses between infancy and childhood

At this stage, we consider two factors potentially playing an important role in the bilateralization of the face-selective response between infancy and preschool childhood. First, infants have poor visual acuity and their visual cortex is therefore exposed mainly to low spatial frequency visual information (Boothe, Dobson, & Teller, 1985; Maurer & Lewis, 2001). A long standing perspective points to a right hemisphere advantage in processing low spatial frequencies of visual inputs (Sergent, 1982; see also Ivry & Robertson, 1998). Even though clear-cut evidence is still lacking, this factor is thought to play a key role in the dominance of the right hemisphere for face perception (Sergent, 1985; Sergent, 1982). According to de Schonen and Mathivet (de Schonen & Mathivet, 1989)’s hypothesis, the right hemisphere may take precedence due to its faster maturation rate at the time at which the first facial inputs, based on low spatial frequencies, are presented to the infant brain. Since the corpus callosum, the main interhemispheric pathway, is not functional before the age of two years (de Schonen & Bry, 1987; Le Grand et al., 2003; Liegeois, Bentejac, & De Schonen, 2000; Salamy, 1978), visual input to each eye is directed only to the contralateral hemisphere in young infants. Thus, it may be that the initial selective representation of faces develops essentially in the right hemisphere, and then transfers to the left hemisphere only after a few years of age, with the maturation of the corpus callosum. Such a developmental scenario would account for the difference between the right lateralization of the response observed between 4-6 months old infants (de Heering & Rossion, 2015) and the bilateral response in preschoolers (here).

A second, non-exclusive potential factor is that although young infants are able to perform perceptual object categorizations (Quinn, Eimas, & Rosenkrantz, 1993; Rakison & Oakes, 2003), they did not yet, arguably, encounter many different categories of objects in their environment. Hence, since it contrasts familiar visual patterns (i.e., faces) with unfamiliar patterns, infants’ right lateralization of the face-selective response observed with this paradigm might be particularly enhanced. However, in 5-year-old children as tested here, stimulation sequences contrast two familiar categories, i.e. faces vs. a wide range of objects (e.g., flowers, fruits, lamps, guitar,…), a factor which may contribute to their lack of RH lateralization. Yet, against this hypothesis, we note that in adults at least, the contrast of faces to non-face objects results in more focal and greater right lateralization in fMRI than the contrast of faces to meaningless patterns such as scrambled faces (e.g., Rossion et al., 2012; Wiggett & Downing, 2008).

### Beyond lateralization: quantitative and qualitative response increase between infancy and childhood

Interestingly, the selective response to faces in preschoolers is of much greater amplitude than in infants, but also much more complex: while it is essentially limited to the first harmonic at 1.2Hz in infants (Figure 1C, (de Heering & Rossion, 2015), the response spreads over multiple harmonics in preschool children (Figure 2). Moreover, the peak and distribution of the response on the scalp also differs, with a relatively focal peak at (right) occipito-temporal sites in infants (P8 channel), as well as in adults (Retter & Rossion, 2016; Rossion et al., 2015), while the bilateral face-selective response observed in children is much more widely distributed, even peaking more dorsally on parieto-occipital sites (Figure 2B).

An increase of the amplitude of the face categorization response is expected between infancy and childhood due to the maturation of the face (and object) processing systems. However, admittedly, we cannot fully exclude an increase in the magnitude of the response also due to the different lengths of stimulation sequences used (i.e., to avoid tiredness, infants were tested with shorter sequences, but were presented with more sequences on average) and a higher attention to the stimuli in children who, contrary to infants, had to perform an explicit orthogonal task to maintain fixation.

At present, we can only conjecture about the factors leading to these qualitative changes between infancy and childhood in generic face categorization. On the one hand, the increase in the number of significant harmonics in the response can be interpreted as an increase in the complexity of the response. While in infants the response appears to mainly reflect a single, low frequency (i.e., 1.2 Hz) face-selective deviation from the 6 Hz base rate used in the stimulation sequence, the children’s face-selective response is likely to be constituted of several differential (i.e. face-selective) components, as in adults (Retter & Rossion, 2016; Rossion et al., 2015). Such a detailed investigation is beyond the scope of the present report. On the other hand, since cortical folding or gyrification is stable despite changes in brain size between infancy and childhood (Armstrong, Schleicher, Omran, Curtis, & Zilles, 1995), the wider spread of the face-selective response over medial occipital and parietal sites in children, independently of its left lateralization, is unlikely to be due to changes in the orientation of the cortical sources with respect to the scalp. Rather, this effect may also be partly related to changes in corpus callosum maturation

### The link between reading acquisition and face categorization

Our EEG data suggest that in 5-year-old children the two hemispheres do not differ in neural specialization for face stimuli. This lack of lateralization cannot be due to a lack of sensitivity of the technique, since the exact same approach reveals a right lateralization in infants (de Heering & Rossion, 2015)and in the majority of typical adults tested (Liu-Shuang et al., 2016; Retter & Rossion, 2016; Rossion et al., 2015). This finding is in line with the lack of significantly larger face-evoked N170 in the RH than in the LH in children up to 9-12 year old, even though the pattern of right lateralization is not systematic across ERP studies (Dundas et al., 2014; Itier & Taylor, 2004b; Kuefner et al., 2010; Taylor et al., 1999). However, here we did not only measure the *absolute* response to faces, but a high-level face-selective response, where faces are contrasted to many variable categories of stimuli. Moreover, the children tested here are younger than the children tested with faces in previous EEG studies with faces and, importantly, they already show a left lateralized response to alphabetic material as a group (Lochy et al., 2016).

Interestingly, across the population of children tested here, there was no significant correlation between this left lateralization of the response to letters and the right lateralization of the response to faces. Although this result does not agree with the correlation between the amplitude of the response to words in the LH and that to faces in the RH in 7-12-year-old children (Dundas et al., 2014), we note that the reverse correlation has also been reported (Li et al., 2013), and that other studies also failed to find significant correlations between the two responses (in adults: fMRI, (Davies-thompson, Tashakkor, & Barton, 2016; Pinel et al., 2014). Moreover, there was no relationship between word and face lateralization as assessed by behavioral performance for hemifield displays in older children and teenagers (Dundas et al., 2012).

Nevertheless, our data show a positive correlation between the pre-reading performance of children and the lateralization of the response to faces: the more letters known by the preschoolers, the more right-lateralized their face categorization response. This is an interesting finding, which supports the view that the right lateralization of face perception is somewhat linked to reading acquisition and competence (Behrmann & Plaut, 2013; Dehaene et al., 2015; Dundas et al., 2012). Note that this result should also be considered in caution, not only because the correlation was not very high, but also in regard of the literature. Indeed, the few studies that tested the relationship between reading abilities and face processing have provided mixed results. In 4 years old tested in fMRI (Cantlon, Pinel, Dehaene, & Pelphrey, 2011), no correlation was found between performance on a face task and right hemispheric response to faces, and between symbols reading and activation for symbols in the same hemisphere. There were however negative correlations between these tasks and non-preferred categories (face task and bold response to shoes in the right; symbols naming and bold response to face *in the left hemisphere*), interpreted as showing that the specialization of these regions for categories does not imply an *increase* of the response to the category but a *decrease* to other categories, which would be due to pruning. In another study comparing several age groups (7-9; 11-13; and adults), the emergence of lateralization to faces (assessed with behavioral hemifield accuracy rates) correlated, as here, with reading competence (Dundas et al., 2012). Yet, the reverse relationship has also been found: higher reading scores associated with reduced face-evoked N170 right-lateralization (Li et al., 2013). Finally, other studies did not measure reading performance, which could therefore not be related to lateralization of neural responses to faces (Dundas et al., 2014).

In summary, it is fair to say that the factors leading to the right hemispheric specialization for faces in the human adult brain remain largely unclear at present, and cannot be simply directly related to reading acquisition: early right lateralization for faces (and face-specific responses) in infants does not support this view, and the relationship between the lateralization of visual language processes and face processes in children and adults remain controversial. A major obstacle to resolve this issue is that different studies carried out in different populations rely on fundamentally different paradigms and behavioral or neural measures, not only in terms of the techniques used but also in terms of the type of contrast performed (i.e., absolute response to faces or differential response to faces vs. control stimuli; absolute response in one hemisphere or lateralization index). In this context, our study contributes to the literature by clearly and objectively isolating in preschool children the same robust generic face categorization (i.e., face-selective) response with the same fast visual periodic stimulation paradigm tested previously in 4-6 months infants and adults. This approach reveals a bilateral generic face-categorization response pointing to a non-linearity of the right hemispheric dominance for generic face categorization across development.

In addition to the great advantage of the paradigm in terms of applicability to different populations, we should emphasize that it provides objective (i.e., frequency-defined) and highly sensitive responses, often at the level of single participants, and does not require to subtract amplitudes obtained in different conditions, as the measured EEG response in an inherent index of generic categorization for faces. Given these advantages, the FPVS-EEG approach should truly open new perspectives in the future to understand the human development of face categorization (Hoehl, 2014) and of the lateralization of brain function in general.

## Acknowledgements

This work was supported by a PAI/UIAP grant PAI/33, an ERC grant (facessvep 284025) and the Belgian National Fund for Scientific Research (FNRS). We also thank the schools and children for their participation.

